# Implications of drift and rapid evolution on negative niche construction

**DOI:** 10.1101/2021.04.26.441094

**Authors:** Phuong Linh Nguyen, Manon Costa, Nicolas Loeuille

**Author notes:** **Correspondence** Phuong Linh Nguyen. **Author contribution** MC, and NL conceived the idea of the project. PLN contributed to the main design of the model, MC and NL contributed to further developments of the model, and provided additional suggestions and modifications. PLN analysed the model with the help of MC and NL. PLN wrote the first draft of the manuscript. MC and NL revised, commented and made further modifications.

## Abstract

1. Organisms throughout their lives constantly modify their surrounding environment; such activities are often termed niche construction. An important property of niche construction is that its consequences can persist for a long period of time and several subsequent generations can be affected. This phenomenon is described as a time lag in niche construction, or ecological inheritance.
2. Studies have suggested that time lag in niche construction can help avoiding the tragedy of the commons. In other words, it can lead to evolution of contribution to a common good, which is associated with positive niche construction, or to the limitation of a common bad, which is associated with negative niche construction.
3. In this article, we will study the evolutionary consequences of incorporating time lags in a negative niche construction process: waste production. We consider a population that extrudes waste into its environment as it consumes resources to grow and reproduce. Higher consumption rates can lead to higher waste production. Individuals that adopt this selfish strategy are expected to be selected as toxic effects are equally shared among all individuals.
4. We show that indeed this tragedy of the commons persists in many cases and selfish strategies evolve in general. When evolution is rapid and intragenerational time lag is incorporated, however, selfish strategies are no longer favoured and strategies resulting in less waste production can be selected. Importantly, heavy pollution results in smaller population sizes, so that drift becomes more important than natural selection and limits the evolution of higher waste production.

## 1 Introduction

Niche construction is a process whereby organisms modify their surrounding environment. It can be as sophisticated and noticeable to the human eye as beaver dams or termite mounds (Naiman et al. 1988; Wright et al. 2002; Korb 2011). Yet, it can simply be a change in chemical concentrations induced by the activities of organisms such as the enrichment of environmental oxygene by cyanobacteria billions of years ago (Mazard et al. 2016). In fact, any living being is a niche constructor because by merely existing, organisms interact with their surrounding environment, thereby chemically and physically modifying it. Such modifications can be positive (positive niche construction) or negative (negative niche construction) when considering the fitness of individuals of the constructing species. It is suggested that niche construction have important ecological and evolutionary consequences (Odling-Smee et al. 2003).

An important property of niche construction is that environmental modifications can persist on long timescales, which is often known as legacy effects or ecological inheritance (Cuddington 2011; Odling-Smee et al. 2003; Danchin et al. 2011; Hastings et al. 2007). In particular, changes in the niche can be inherited within a generation and between generations of a niche constructing species (Krebs and Davies 1993; Laland et al. 2000; Edeline et al. 2016; Hastings et al. 2007). Environmental changes can also be inherited by other species that live within the same area (Hastings et al. 2007; Kidwell and Jablonski 1983).

Understanding the evolutionary dynamics related to niche construction therefore requires the careful consideration of three different timescales: the population timescale, the niche construction timescale, and the evolutionary timescale. The population dynamic timescale encompasses all demographic processes of niche constructors and recipients of niche construction. The niche construction timescale covers the variations in the environment born from niche construction processes, including ecological inheritance. Finally, the evolutionary timescale refers to the changes in gene frequencies, emergence and invasion of new mutants, or the birth and death of new species. For instance, a termite mound may grow as the termite colony grows; this happens along the population dynamic timescale. The changes of the mound could then affect local environments for millennia (Martin et al. 2018), so that the niche construction timescale here is very large. Associated environmental changes can have large consequences, affecting vegetation patterns at various spatial scales (Bonachela et al. 2015; Tarnita et al. 2017; Ashton et al. 2019) thereby creating new sources of selection that act on a long evolutionary timescale.

The three timescales thus interact in complex ways and do not necessarily match. If niche construction persists for a long time, its timescale may completely lag behind the population dynamic timescale. For instance, mollusca or crustacean species leave behind their shells when dead, which accumulate under the ocean. This gradually forms hard substrata which facilitate or inhibit the occupation of subsequent species (Kidwell and Jablonski 1983). In this case, several populations may exist, reach their dynamical equilibrium, and even go extinct, while the dynamics of the substrata remains at its quasi-stable state. Thus, the substrata dynamics may not have a significant effect on a particular species within a short period of time, but when considering a sufficiently long period, the effect becomes more significant and concerns evolution of multiple species. The lag between the population and niche construction timescales need not be so extreme (Odling-Smee et al. 2003). For instance, earthworms modify soil properties which has been suggested to make the environment become favourable for not only the starting communities, but also their future generations (Caro et al. 2014). In the study of Edeline et al. (2016), when juvenile and adult mekada fishes consume the same resources, adults ‘inherit’ the resources degraded by juveniles, and this facilitates the evolution of semelparity.

Theoretical frameworks of evolution often assume the separation of the evolutionary and population dynamic timescales, such that the former is much slower than the latter (Metz et al. 1995; Koch et al. 2014). However, more and more evidence pointed out that evolutionary processes can be much faster than previously thought (Thompson 1998; Hairston et al. 2005; Carroll et al. 2014), so that evolutionary and ecological timescales may not be easily separated. If the dynamics of niche construction are also taken into account, then lags among the three timescales can happen in many ways, leading to unexpected ecological and evolutionary results. For instance, Gurney and Lawton (1996) showed that a time lag between population and resources dynamics, can lead to cyclic dynamics. Laland et al. (1996) showed that a lag in the effect of resources construction delays the spread of the allele that is favoured by the increasing amount of resources. Both studies concern positive niche construction where a focal population can increase resources dynamics.

Effects of niche construction, positive or negative, are often shared among coexisting individuals and may result in the tragedy of the commons. It is often difficult for positive niche construction to evolve but easy for negative niche construction to spread. To avoid this tragedy, classical theoretical studies include a direct benefit to the restriction of negative niche construction or impose a direct cost by coercion and punishment, or add spatial structure and kinship (Rankin et al. 2007). They have one thing in common: explicit dynamics of niche construction are not taken into account, that is, organisms can impact the environment but feedback loops between the environment and organisms are disregarded. Such feedback loops are however suggested to change evolutionary dynamics (Odling-Smee et al. 2003; Estrela et al. 2019).

In this article, we explicitly include all three dynamics: population, niche construction and evolution and consider possible lags among the three associated timescales. We study the evolution of negative niche construction, here the production of waste. Waste production is assumed to be positively linked to consumption rates such that individuals that consume more produce more waste (Zarco-Perello et al. 2019; Besiktepe and Dam 2002; Tanner et al. 2019), and have higher reproduction, growth or maturation rates (Greenberg et al. 2003; Morton 1986). However, waste production also pollutes the environment, thereby reducing the fitness of the population. When such fitness reductions lead to smaller population sizes, they may increase the significance of genetic drift compared to natural selection. We found that in almost all cases, negative niche construction is favoured, possibly leading to population extinction. To counterselect for it, we need to introduce intragenerational time lags between niche construction and population dynamics. Also, evolutionary timescale needs to be overlapped with the other two timescales. More importantly, since negative niche construction leads to small population sizes, drift plays an increasing role compared to natural selection which may limit negative niche construction activities.

## 2 Model

The analysis is structured as follows: we first use the adaptive dynamics approach to analyse scenarios of slow evolutionary dynamics (Metz et al. 1995). The most important assumption of this approach is that the evolutionary timescale completely lags behind the other two timescales. Therefore, mutation is so rare that when a new mutant arises, the resident population is already at its ecological equilibrium, setting the environmental conditions, and the mutant will replace the resident if its invasion fitness is positive. We then incorporate intragenerational time lags using a structured population model, where the population is divided into juvenile and adult states. Here, the intragenerational time lag implies that adults are affected by the environment constructed by juveniles. Negative niche construction thereby directly affects individual fitness. Finally, we relax the assumption of slow evolution imposed by the adaptive dynamics approach. Multiple mutants can arise at the same time when resident populations need not be at the equilibrium. As a result, offspring with different strategies inherit the environment created by previous generations. We use the Tau-leap method to simulate the dynamics (Gillespie 2001). This method enables an overlap between the evolutionary timescale and the population and niche construction timescales. We will denote this overlap as rapid evolution. This also allows us to study the effect of drift because birth and death processes are modelled as stochastic drawings. As negative niche construction can lead to smaller population sizes, the role of drift can become more significant.

### 2.1 A complete lag of the evolutionary timescale

#### Negative niche construction without intragenerational time lag

We model a species *S* that impoverishes its environment by consuming resources *R* and pollutes it by producing waste *W*. A higher consumption rate *c* results in more offspring, given a fixed efficiency *ρ* of converting resources into new individuals. It also leads to higher rates of waste production *f*(*c*), where *df*(*c*)*/dc >* 0. The pollution level (*W* adds a mortality rate *ω*(*W*) to the natural mortality rate *d* of the consumer. Since higher waste densities lead to higher additional mortality rates, we assume *dω*(*W*)*/dW >* 0. The dynamics of resources and waste follow a chemostat dynamic, where *I*_*R*_/*δ*_*R*_ and *I*_*W*_/*δ*_*W*_ are their respective natural turnover rates. The ODEs that describe the whole system can be written as

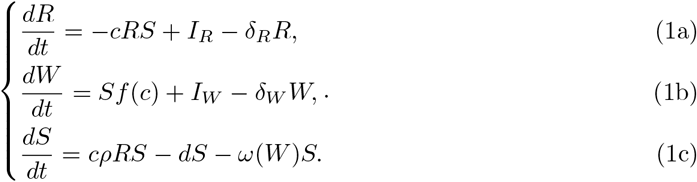

In order to derive analytical results, we use linear functions for the production of waste and additional mortality due to pollution, thus, *f*(*c*) = *hc* and *ω*(*W*) = *vW*. System (1) has three equilibria: an equilibrium where the species does not survive, an equilibrium where the density of the species is always negative, and an equilibrium where the species persists if the consumption rate is sufficiently large, i.e. a positive equilibrium. This positive equilibrium is unstable only if the niche construction activity has little effect on the waste dynamics or if the population is not at all vulnerable to pollution. This results in the population increasing to infinity, and it corresponds to extremely small values of *h* and *v* (details of the equilibrium is in Supplementary Document 1). In our analysis, we only consider sufficiently large values of *h* and *v*, such that the equilibrium is always stable.

We study the evolution of consumption rate *c*. A mutant that adopts a different consumption value than the resident can invade if its invasion fitness is positive. This is the equivalent to a mutant having its reproduction ratio *F*_*m*_(*c*_*m*_, *c*) greater than one. In other words, a mutant can spread if it is replaced by more than one offspring (details of the expression is in Supplementary Document 2). The reproduction ratio of a mutant, which is derived from the invasion fitness condition, can be written as

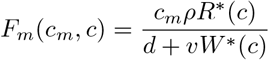

The value of the reproduction ratio depends on the mutant strategy (*c*_*m*_) and the environment that is constructed by the resident, which is evaluated at equilibrium (*W*\*(*c*)*, R*\*(*c*)). It can be shown that the selection gradient on higher consumption rate is always positive (see details in Supplementary Document 2). As a consequence, we always observe selection for higher consumption rates, leading to a continuous increase in pollution and more scarcity of resources (figure 1). The consumer population eventually settles to a certain value, when increasing consumption rates are exactly balanced by increased costs due to pollution. Note that the selection pressure remains positive, but its value decreases as the consumption rate increases (purple line in figure 1), so that evolution becomes progressively slower.

**Figure 1:**
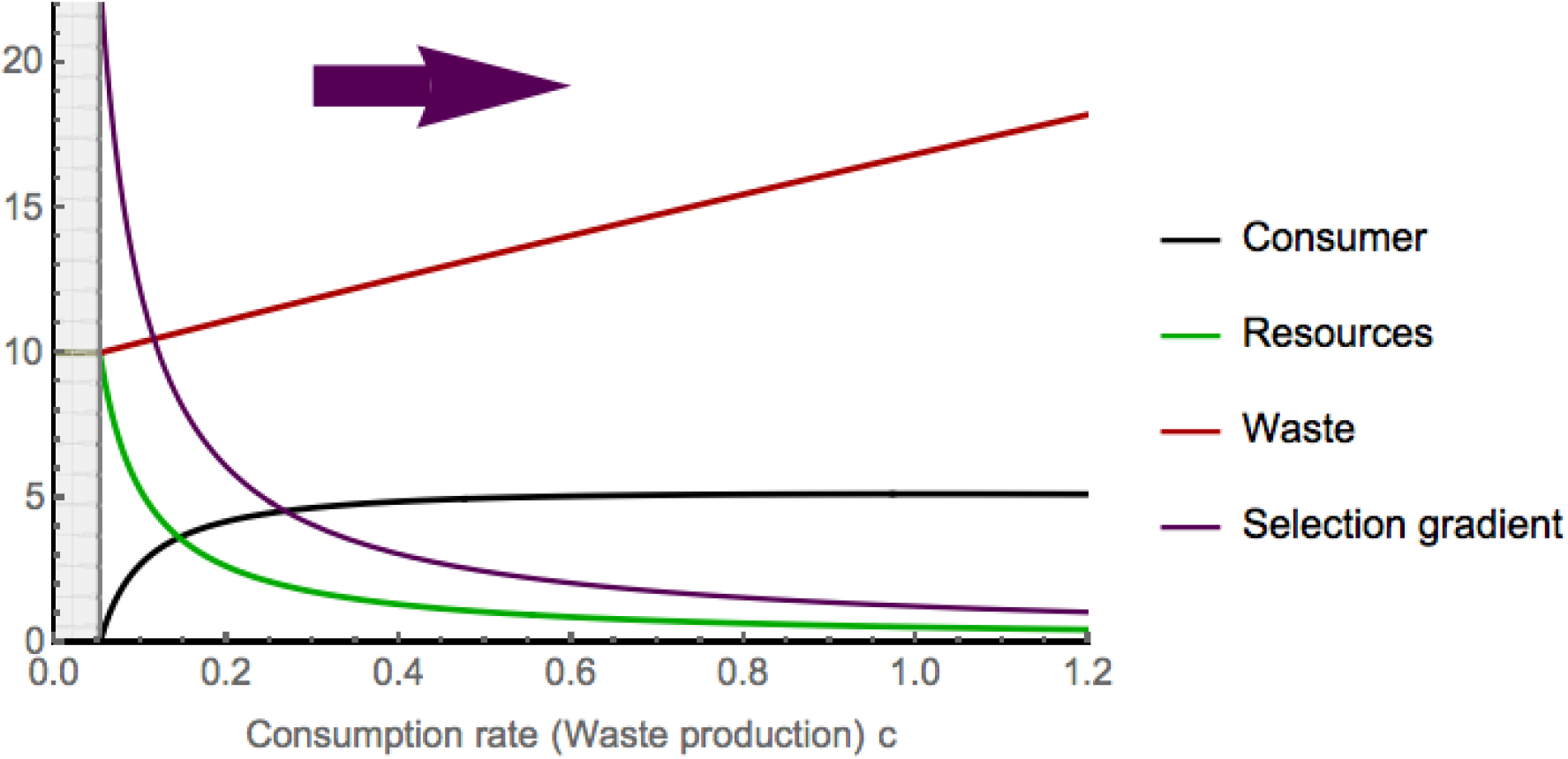
Changes in equilibrium value (*W*\*(*c*), *R*\*(*c*), *S*\*(*c*)) with respect to the trait value. Gray area corresponds to population extinction. Parameters: *ρ* = 2.3*,d* = 1.1, *I*_*R*_ = 3, *δ*_*R*_ = 0.3, *I*_*W*_ = 3, *δ*_*W*_ = 0.3*,v* = 0.01*,h* = 0.4

In this model, there is no direct cost on over-exploitation, and the tragedy of the commons persists. All individuals, consumptive or frugal, share the damage caused by high pollution levels and resource degradation but the benefits to reproduction is attributed immediately to the individuals that adopt the selfish strategy of overexploitation.

#### Negative niche construction with intragenerational time lag

Intragenerational time lag is incorporated using an age-structured population, in which a consumer has a juvenile state (*J*) and an adult state (*A*). Juveniles mature into adults at a rate *g* and adults reproduce at a rate *ρ*. Both the maturation rate and the reproduction rate are functions of juvenile and adult consumption rates (*c*_*J*_, *c*_*A*_) and of resources availability (*R*). In consuming resources, both juveniles and adults emit waste at a rate *p*_*J*_(*c*_*J*_) and *p*_*A*_(*c*_*A*_) respectively. These rates depend on the consumption rates of adults and juveniles, such that the more they consume, the more they pollute their environment. Juveniles and adults have an additional mortality rate due to pollution (*ω*_*J*_(*W*) and *ω*_*A*_(*W*)). The intragenerational time lag in niche construction is depicted by the fact that the negative effect of resource degradation and pollution is transmitted from juvenile to adult states. The natural mortality rates of juveniles and adults are *d*_*J*_ and *d*_*A*_ respectively. As before, the dynamics of the resources and waste follow a chemostat dynamic. Their natural turnover rates are respectively *I*_*R*_/*δ*_*R*_ and *I*_*W*_ /*δ*_*W*_. The ODEs that describe the dynamics of the system read

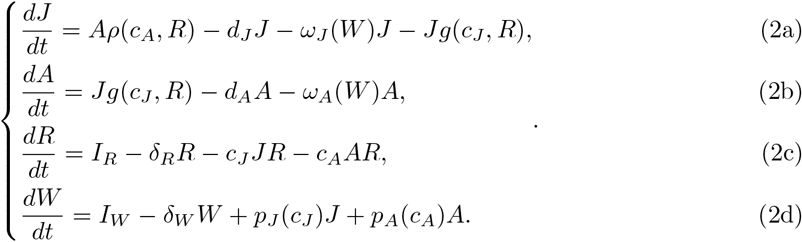

System (2) is rather complicated to analyse theoretically. The number of equilibria depends much on the explicit forms of the additional mortality functions (*ω*_*J*_(*W*) and *ω*_*A*_(*W*)), and the reproduction and maturation functions (*ρ*(*c*_*A*_, *R*) and *g*(*c*_*J*_, *A*)). Even when we use all linear functions, it is still difficult to obtain analytical results. This complicates our evolutionary analysis as the invasion fitness of a mutant depends on the value of the resident at equilibrium. Therefore, we simplify the model to gain a better understanding of how the environment affects the selective pressure.

We only consider negative niche construction as increases in pollution levels, disregarding the dynamics (and overexploitation) of resources. In addition, we use linear relationships in all functions. System (2) can now be simplified into

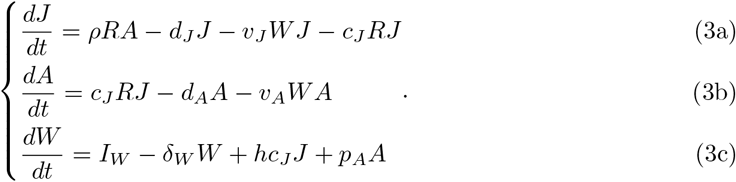

Now, resource density *R*, reproduction rate *ρ* and waste production rate of adults *p*_*A*_ are constants. *v*_*J*_, *v*_*A*_ are the vulnerabilities of juveniles and adults to pollution. In disregarding the resources dynamics and considering exclusively linear functions, the population dynamics are entirely governed by the waste dynamics. There is thus no resource competition among individuals, adult and juvenile alike. Our model becomes similar to models of maturation (Roos et al. 2007; Gardmark et al. 2003; Poos et al. 2011). Very often in these models, there is a trade-off between adult reproduction and juvenile maturation, such that, when an individual invests more in maturation, it invests less in reproduction because it has a fixed energy budget. In our model, considering such a trade-off would correspond to the consideration of an intrinsic constraint of the negative niche construction activity, which, similar to the study of Kylafis and Loreau (2008), may result in selection of lower negative niche construction. In this article, we investigate whether such reductions in negative niche construction may arise only from variations in the different timescales, and therefore do not include direct costs. Therefore, we assume no direct link between juvenile and adult traits.

System (3) has three equilibria: one trivial equilibrium where no adults and juveniles can survive, one equilibrium where the waste density is always negative, and one equilibrium that is positive if *F >* 1, where

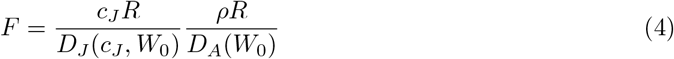

is the reproduction ratio of a resident consumer. Here, 1*/D*_*J*_(*c*_*J*_, *W*_0_) = 1/(*d*_*J*_ +*c*_*J*_*R*+*W*_0_) is the expected time that the consumer spends as juvenile, and 1*/D*_*A*_(*W*_0_) = 1/(*d*_*A*_ + *W*_0_) is the expected time that the consumer spends as adult, with *W*_0_ = *I*_*W*_/*δ*_*W*_ is the waste density in the environment when the consumer is rare. *F* includes both adult and juvenile components, thus, *F >* 1 requires that the adult reproduction rate and the juvenile maturation rate are sufficiently large, while the waste turnover has to be sufficiently small so that the environment is livable. When *F >* 1, the equilibrium is most likely stable (details of the equilibrium are in Supplementary Document 3 and 4).

We analyse the evolution of the juvenile consumption rate. A lower consumption rate indicates lower rates of maturation and waste production. A mutant with a consumption rate *c*_*Jm*_ can invade a resident population whose dynamics are at equilibrium if its invasion fitness is positive. This condition is satisfied whenever the mutant reproduction ratio *F*_*m*_ is greater than one (details in 5), where

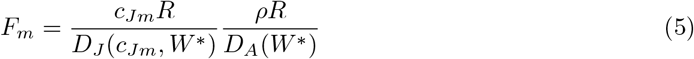

Here, 1*/D*_*J*_(*c*_*Jm*_,*W*\*) = 1/(*d*_*J*_ + *c*_*JmR*_*R* + *v*_*J*_*W*\*) is the expected time the mutant spends as a juvenile, and 1*/D*_*A*_(*W*\*) = 1/(*d*_*A*_ + *v*_*A*_*W*\*) is the expected time the mutant spends as an adult. *W*\* is the waste density at equilibrium, which depends on the growth rate value *c*_*J*_ of the resident. The reproduction ratio of a mutant (*F*_*m*_) is rather similar to the reproduction ratio of a resident (*F*), except that the latter depends on a ‘virgin’ environment whereas the former depends on the environment constructed by a resident. Expression (5) suggests that higher juvenile consumption reduces the time that a consumer spends as an adult because it increases the pollution level so that the consumer might die before it can even reproduce. Thus, a lower juvenile consumption might be selected under certain conditions. This happens when the selection gradient is negative, which requires

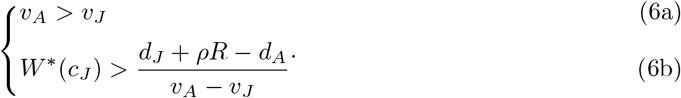

Condition (6) suggests that the direction of the selection gradient does not depend on the mutant trait value, and essentially depends on the pollution level created by the resident. Condition (6a) implies that adults have to be more vulnerable to pollution than juveniles. Intuitively, if juveniles are more prone to pollution than adults, those who mature slower remain juvenile for a longer time and suffer pollution, whereas those who mature faster escape the (vulnerable) juvenile state. Selection then always favours higher juvenile consumption. Thus, in order for lower trait values to be selected, adults have to be more vulnerable to pollution than juveniles. The second condition (6b) implies that if the waste density at equilibrium is sufficiently large, the environment becomes too toxic, and traits that reduce pollution levels (i.e. lower consumption) may be selected.

However, under the assumptions of adaptive dynamics, condition (6b) can never be satisfied. When the environment is not too polluted, we observe a strong selection pressure for higher consumption rate (figure 2A, S. 2A). As the consumption rate increases, so does waste density (figure 2A) (see Supplementary Document 6 for more details). Because mutants can only arise when the resident population is at equilibrium, and because the waste density at equilibrium is asymptotic to (*ρR - d*_*A*_)*/v*_*A*_ as the evolving trait increases, which is smaller than the threshold set by the right-hand side of condition (6b) (figure 2, Figure S. 2), higher consumption is always favoured. This selection leads to a continuous increase the pollution level (Figure S. 3), and so the tragedy of the commons persists. Note, however, that the selection gradient fastly becomes extremely small, so that we expect selection for higher consumption rates to become very weak, and evolutionary dynamics to be very slow.

**Figure 2:**
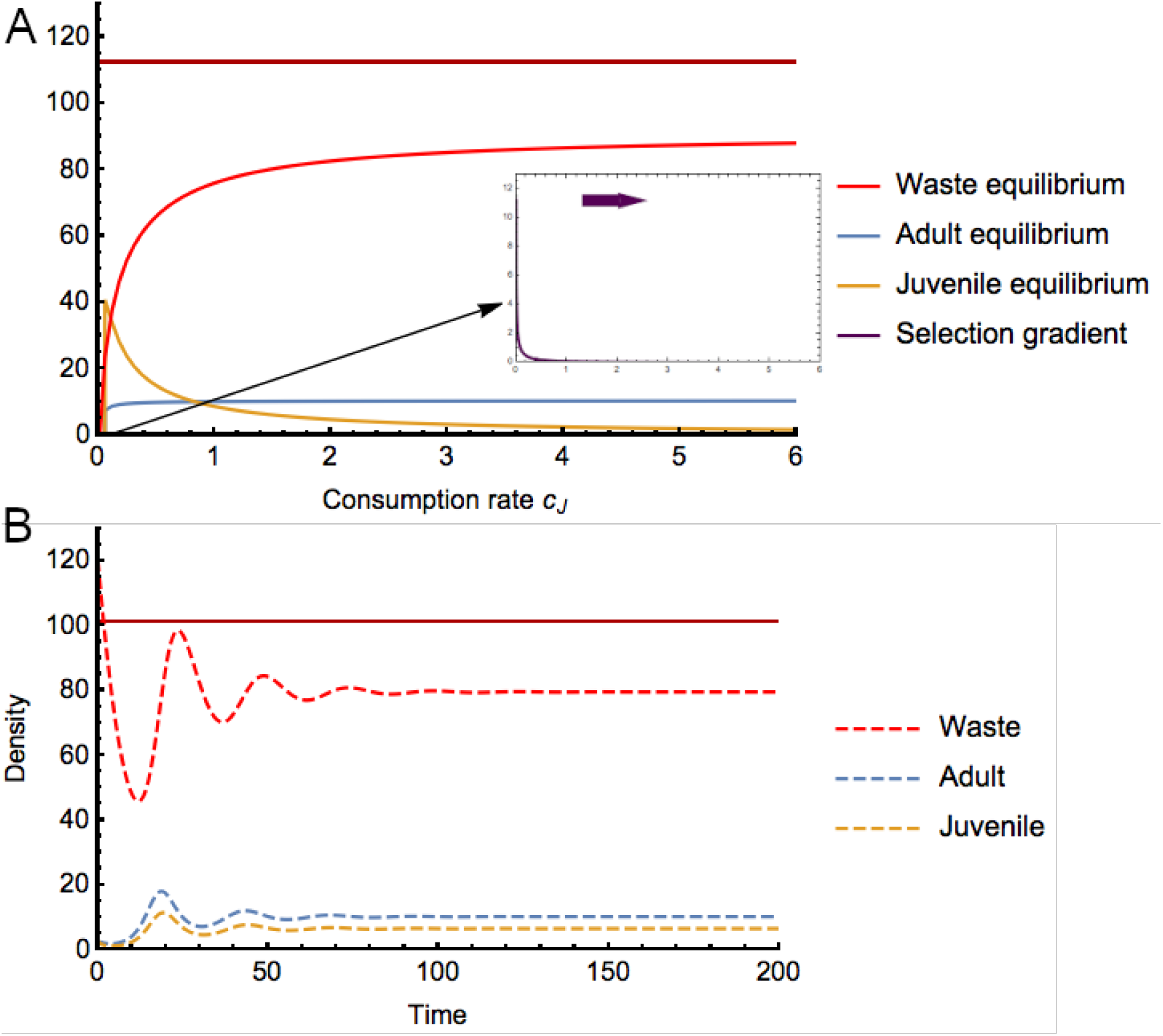
A) Changes of the equilibrium values of system (3) with respect to the growth rate values. The small frame illustrates the selection gradient, and corresponds to a zoom of the general figure B) Ecological dynamics of waste and a resident population that adopts a growth rate value *c*_*J*_ = 1.4. Thick lines indicate equilibrium while dashed lines indicate the value of density of a resident population. The dark red line indicates the threshold beyond which lower growth rate can be selected. Other parameters: *R* = 1, *v*_*J*_ = 0.001, *d*_*J*_ = *d*_*A*_ = 0.1, *h*_*J*_ = 1.1, *v*_*A*_ = 0.01, *ρ* = 1.01, *I*_*W*_ = 0.3, *δ*_*W*_ = 0.13, *pA* = 0.001

### 2.2 Rapid evolution and the role of drift

From condition (6), we infer that if, out of equilibrium, the waste density exceeds its equilibrium value, it can satisfy condition (6b), resulting in selection for mutants with lower trait values. In addition, when the environment is polluted, population density decreases and the selection pressure is weakened (figure 2A), suggesting that drift can play an increasingly dominant role in the evolutionary dynamics. To investigate these aspects, we relax the assumption of slow evolution and introduce stochasticity using mathematical simulations.

In particular, we use the Tau-leap algorithm (Gillespie 2001). At each interval *τ*, we calculate all rates for maturation, reproduction, mortality of juveniles and adults, and the influx and outflux of the waste concentration. Changes in the number of juveniles and adults and in waste concentration are then drawn from a Poisson distribution depending on their respective rates. An increase in the number of juveniles implies birth events. Mutations can happen at a certain rate *m*, and new mutants will adopt juvenile consumptions that are drawn from a normal distribution whose mean is the value of the mother and the standard deviation is *σ*. When *m* is extremely small, we recover the adaptive dynamics scenarios. Rapid evolution takes place when we increase the mutation rate. It should be noted that here rapid evolution implies overlaps between the three timescales and not indicates larger standing variation or stronger selection as in Koch et al. 2014 and Hairston et al. 2005. In fact, the evolutionary speed could vary in the simulations.

In contrast with the adaptive dynamics approach where the population dynamics are deterministic and small populations are guaranteed to survive as long as they satisfy survival conditions, in stochastic simulations, small populations with selective advantages can go extinct whereas those with selective disadvantages can survive simply due to chance, so that the stochastic simulations allows us to consider drift. In each simulation, we start with a monomorphic population and an initial value of waste density that is drawn from a uniform distribution with a range of (1, 10). Such initial values allow the existence of initial populations that are sufficiently large, in an environment that is not too polluted.

#### Negative niche construction with intragenerational time lag

Our simulations suggest that in the long term, an increase in juvenile consumption rate that leads to an increase in waste production is inevitable. However quasistationary states of the trait value are obtained mostly because the low population sizes allow a strong effect of drift, that may easily compensate for the low selection gradient we observed in the adaptive dynamics analysis (figure 2A). Note also that higher trait values are also counterselected whenever the waste density crosses the threshold (*d*_*J*_ + *ρR - d*_*A*_)/(*v*_*A*_ - *v*_*J*_) in condition (6b) (figure 3, 4).

**Figure 3:**
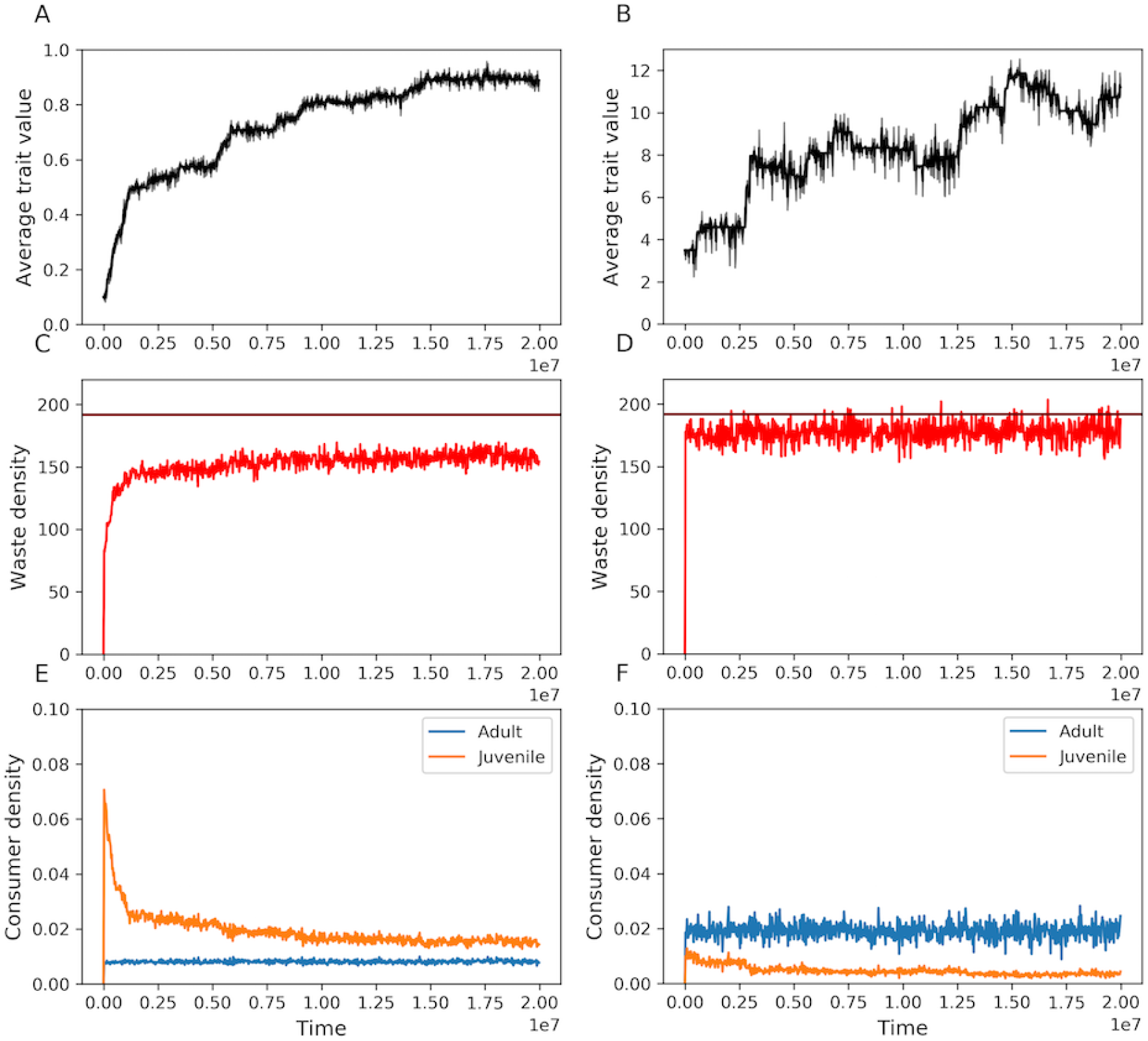
Simulations with moderately fast population and waste dynamics compared to evolutionary dynamics. A, C, E) The starting population has low growth rate *c*_*J*_ = 0.1. B, D, F) The starting population has higher growth rate *c*_*J*_ = 3.5. Other parameters for dynamics of populations and waste: *d*_*J*_ = *d*_*A*_ = 0.1, *h*_*J*_ = 1.1, *v*_*J*_ = 0.0001, *v*_*A*_ = 0.01, *u*_*A*_ = 1, *u*_*J*_ = 1, *I*_*W*_ = 0.3, *δ*_*W*_ = 0.13, *ρ* = 1.9, *p*_*A*_ = 0.001. Parameters for evolutionary dynamics *σ* = 0.02 for low growth rate and *σ* = 0.7 for higher growth rate, *m* = 0.001. Red horizontal lines indicate the threshold for the waste density beyond which selection will favour lower growth rate.

**Figure 4:**
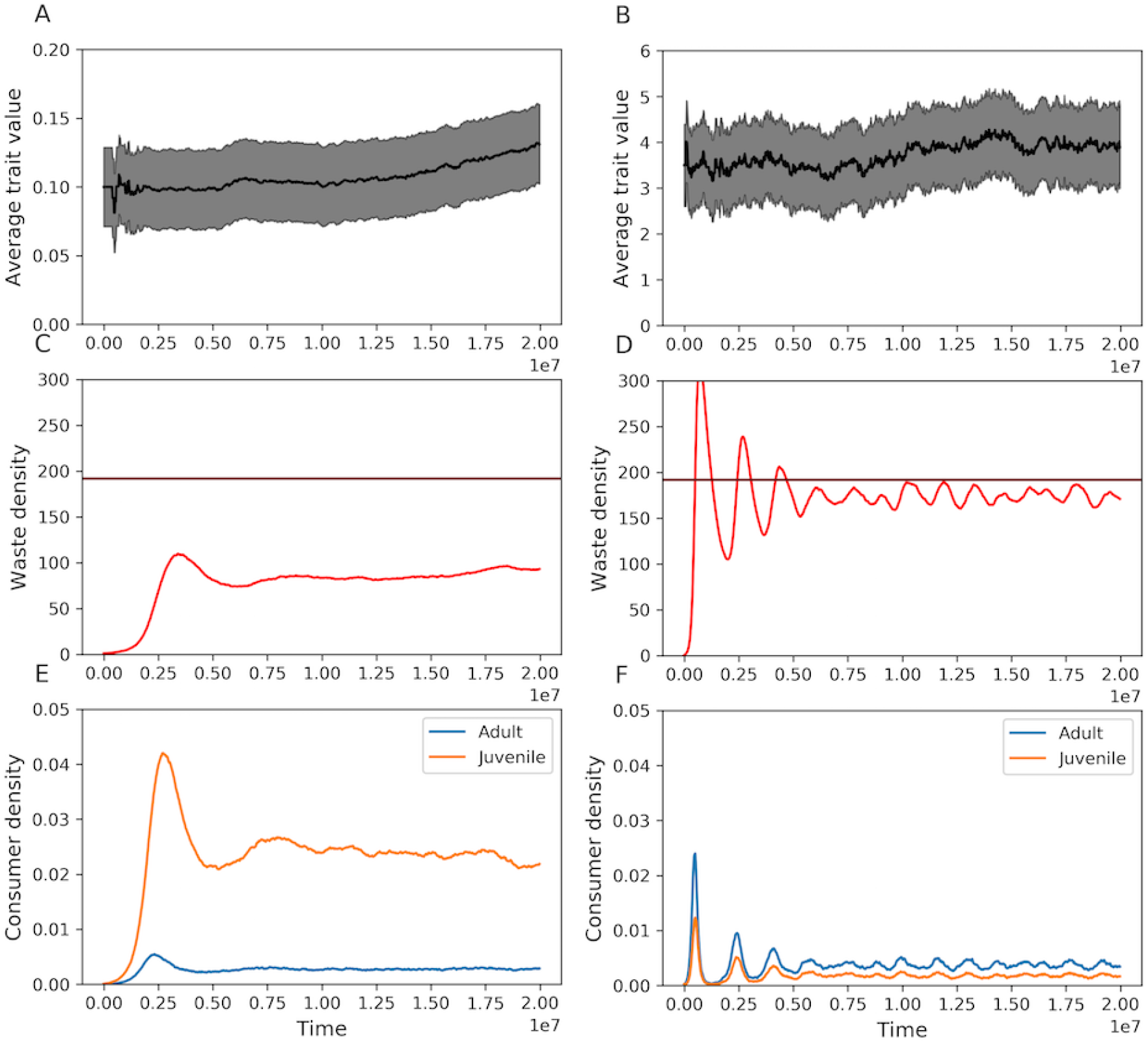
Simulations with slow population and waste dynamics compared to evolutionary dynamics. The dynamics of population and waste are three order of magnitude slower than in figure 3. Mutation rate is increased to *m* = 0.01. Red horizontal lines indicate the threshold for the waste density beyond which selection will favour lower growth rate. The gray area is the standard deviation of the trait value.

When the initial population has an extremely small juvenile consumption rate (*c*_*J*_ = 0.1), higher trait values will be immediately selected because the starting environment is rather clean and the selection pressure is rather strong (left panels of figure 3). This will lead to a rapid increase of waste density. As the environment becomes polluted, the selection pressure for higher consumption is progressively eroded, so that evolution rapidly slows down. When the initial population possesses an already high juvenile consumption rate (*c*_*J*_ = 3.5), the environment becomes instantly heavily polluted and the waste density crosses the threshold. This leads to the counter selection of high trait values. Polluted environments then reduce population density, which subsequently leads to a decrease in the pollution level below the threshold. Higher consumption rates are then again favoured. However, as long as the pollution level is not too far below the threshold, selection pressure for higher consumption rates remains weak while population density is low, so that drift becomes increasingly important. Also, low population sizes lead to few mutations, further limiting evolution toward higher trait values. For all these reasons (limited mutations, weak selection, important drift), we observe that the trait value fluctuates around a quasistationary state that maintains the system close to the pollution threshold (right panels of figure 3).

When the population and waste dynamics are extremely slow whereas evolution remains rapid, a higher juvenile consumption rate is still selected for when the initial population has a small trait value (figure figure 4A). However, the increase in the trait value is much slower than when the population and waste dynamics are fast. When the initial population has a higher trait value, we observe a longer period of quasistationary state of the trait value (figure 4B). In both cases, the variation of the trait values is much higher than when the population and watste dynamics are fast. What is more interesting is that the quasistationary state is initially obtained as a result of counter selection of higher consumption rates because the waste density is above the threshold. However, in the long term, stasis in the trait is maintained mostly by drift, as the waste density remains slightly below the threshold most of the time (right panels of figure 4).

#### Negative niche construction without intragenerational time lag

Our results so far suggest that intragenerational time lag alone is not sufficient to prevent higher waste production. This requires both drift and rapid evolution. In this section, we revisit the unstructured system (1) and analyse whether including drift and rapid evolution is sufficient to avoid higher waste production.

We run simulations adopting the Tau-leap method just as we did for the structured system (3). We found that in all cases, selection for lower consumption rate that is linked to lower waste production, never takes place (figure 5) (more simulations with different parameter values can be found in figure S. 7). The effect of drift is less significant in this case because the population density is large as higher consumption rate is now associated with higher reproduction rate. Selection for higher consumption rate is therefore stronger than the drift effect, and even quasistationary state of the trait value cannot be maintained. This result suggests that rapid evolution and drift do not suffice for the counter selection of negative niche construction, and that an intragenerational effect of niche construction is additionally required.

**Figure 5:**
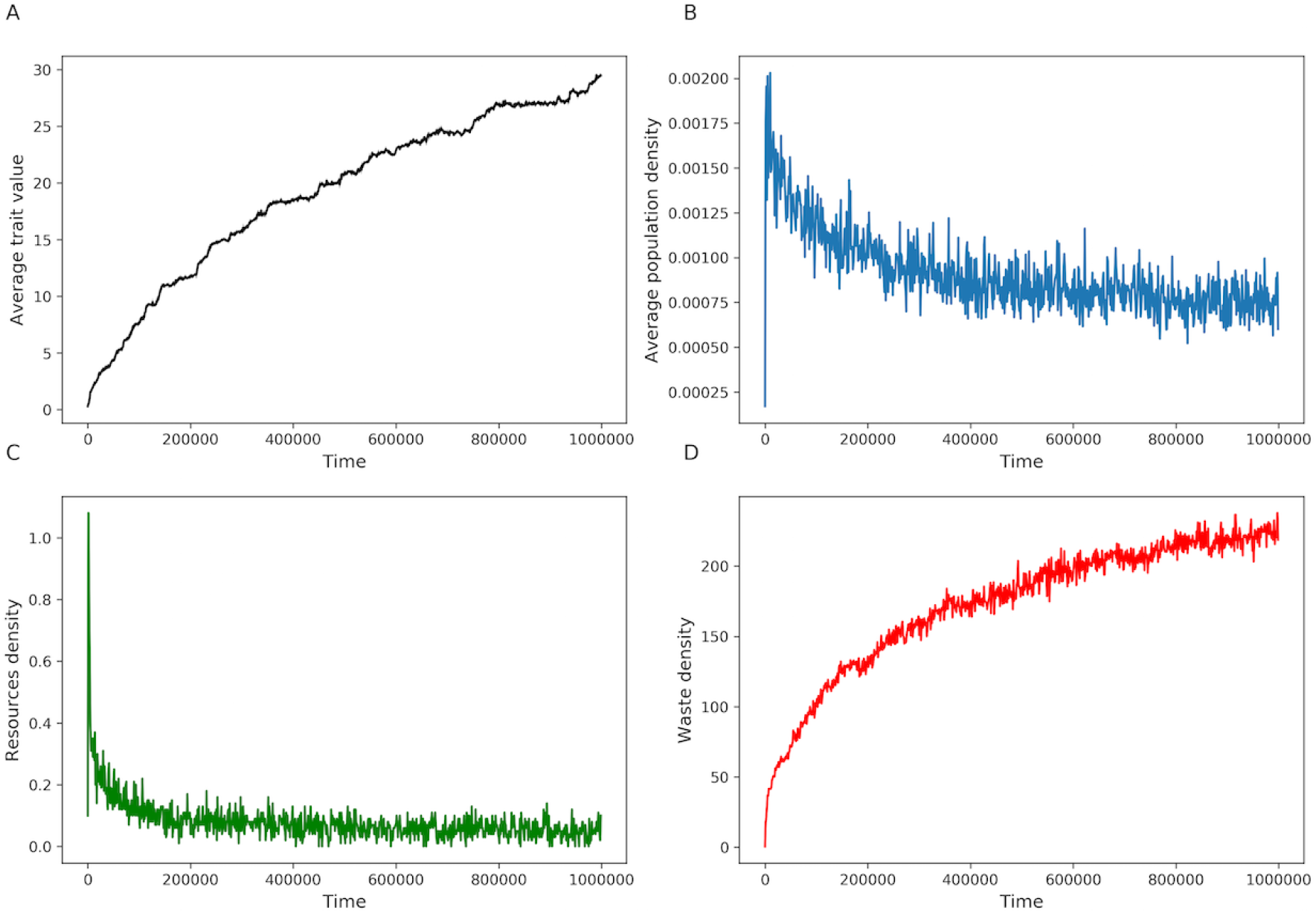
Simulation using Tau-leap method. Parameters: *ρ* = 2.3*,d* = 1.1, *I*_*R*_ = 3, *δ*_*R*_ = 0.3, *I*_*W*_ : 3, *δ*_*W*_ = 0.3*,h* = 1.1*,v* = 0.01*,m* = 0.1, *σ* = 0.2.

## 3 Discussion

Along their life cycles, organisms consume resources and produce metabolic wastes, which inevitably results in resource depletion and pollution. Such environmental modifications may lead to population extinction if the resources become too scarce or the environment too toxic. These poor conditions can persist for a long period of time, and so do their negative effects on the organisms. This is often known as the time lag in niche construction or ecological inheritance (Odling-Smee et al. 2003; Danchin et al. 2011; Cuddington 2011).

In this article, we use mathematical models to study the evolution of negative niche construction manipulating explicitly three different timescales: population, niche construction, and evolution. In addition, because negative niche construction can be associated with decreasing population sizes, we also consider how these small population sizes can affect the evolutionary dynamics. In such conditions, mutations are limited and drift eventually compensates natural selection so that negative niche construction is slowed down. Our results also suggest that intragenerational time lag in niche construction is the precondition but rapid evolution is required for the counter selection of negative niche construction. In addition, drift plays a more important role than natural selection to maintain quasistationary states of the trait value.

Increasing environmental pollution is unavoidable under the adaptive dynamics approach, which assumes that the evolutionary timescale lags far behind the population and niche construction timescale. A mutant with a higher reproduction rate always replaces a resident population despite the fact that it will worsen the environment for both of them. When the environment is heavily polluted, a strain that adopts an overexploitation strategy may die faster but it also reproduces faster to maintain its existence. Eventually, evolution leads to increasing pollution level and decreasing population density, possibly threatening the evolving population. This result has been observed in the study of Ratzke et al. (2018), in which a strain of soil bacteria increases the environmental PH, which in turn becomes toxic to the bacterial population. The bacterial population then collapses quickly because they cannot live in a highly acidic environment.

To prevent such tragedy of commons, direct benefits are usually added to positive niche construction and direct costs are imposed on negative niche construction. For instance, Krakauer et al. (2009) shows that benefits can come from the ability of organisms to monopolise their niches and prevent free riders; Kylafis and Loreau (2010) and Chisholm et al. (2018) suggest that benefits could also be attributed to the ability to better exploit or adapt to the constructed niche. The benefits from positive niche construction in Lehmann (2008) comes from kinship and transgenerational time lag in niche construction.

In the present work, the cost on waste production lies in the intragenerational time lag in niche construction. This potentially creates a threshold of pollution beyond which strains that produce less waste and mature slower have more advantages than strains that mature fast but produce more waste. However, under the adaptive dynamics approach, the waste density always settles at its ecological equilibrium which is below the threshold. Therefore, the advantageous environment for having a slow maturation rate vanishes when mutations emerge. That is why rapid evolution is mandatory, where the evolutionary dynamics can be on a similar timescale as the waste and population dynamics. In such a case, high pollution levels may persist while strains with slow and fast maturation rates coexist. Juveniles who produce more waste mature faster into adults and pay a higher cost. By contrast, juveniles who produce less waste mature slower, remain juvenile for longer and pay a smaller cost. Halting negative niche construction also requires that adults are more vulnerable to pollution than juveniles. Here, the negative effects of pollution are shared among individuals but the costs on different strategies are unequal. In our model, rapid evolution allows rapid feedback loops between evolutionary dynamics, niche construction and population dynamics. It has been shown that such rapid feedback loops play a key role in the evolution of positive niche construction. In the studies of Weitz et al. (2016) and Tilman et al. (2020), reckless consumption cannot prevail. It is beneficial in a nutrient rich environment, and so the frequency of individuals that adopt this strategy will increase. However, along with this increase, they impoverish the environment and the reckless consumption strategy is now at a disadvantage compared to the prudent consumption strategy.

Evidence for rapid evolution is rich. For instance, changes in beak and body size of Darwin’s finches and changes in the diapause timing of a copepod species happen within a few generations (Grant and Grant 1995; Hairston and Dillon 1990). Many more examples can be found in Hairston et al. (2005) and Thompson (1998). Studies on the effect of intragenerational time lag of niche construction are however rare. A study of positive niche construction in *Coenobita compressus*, a terrestrial hermit crab, may give a hint of the importance of intragenerational time lag. *C. compressus* has been shown to be able to modify the shells they reside in (Laidre et al. 2012; Laidre 2012a), and when they outgrow their current shell, they change to a bigger shell. Laidre (2012b) shows that the crabs prefer modified shells that have been used by other crabs because the modified shells increase their survivorship. The used shells that they abandoned will serve as new shells for other smaller and younger crabs. Here, “juveniles” are affected by positive niche construction activities of “adults”. As generations overlap, such modifications would still be considered as intragenerational time lags in our population structured framework. Importantly, juveniles and adults involved in this example do not have to be kin.

One important result is that in the long term, drift plays a key role in preventing the increase of waste production. Early rapid evolution leads to the selection of highly consumptive traits that lead to a heavily polluted environment. As the waste density may temporarily reach high values (above the threshold), strains that produce less waste can become temporarily advantageous. This then results in smaller population density and a less polluted environment in which strains that produce more waste and mature faster again have more advantage. However, as the environment becomes progressively occupied by many strains, the pollution level remains high. This situation has two immediate consequences: (i) population density is kept at a low value, and (ii) the selective force favouring higher waste production becomes very small. Drift then becomes dominant and evolutionary trajectories fluctuate without a clear direction (quasi stationary state). It should be noted that the effect of drift is specifically important in our model on negative niche construction because negative niche construction may lead to decreasing population size. We expect that the drift effects we observe may not be that important if niche construction is positive because positive niche construction by definition leads to higher fitness within the population which may often (but not always) lead to higher population sizes. Such higher population sizes should favour the action of natural selection over drift.

In our intragenerational model, we exclude the effect of resources availability on the selection pressure, which could be an important component to prevent the increase of waste production. In fact, Kawecki (1993) showed that if there is competition for resources among juveniles and adults, individuals that delay maturation may grow larger, obtain more resources and therefore produce more offspring than individuals that mature early. Future studies that take into account resources dynamics would provide a deeper understanding.

Our models are simple but they take into account two most fundamental elements: a niche constructing population and the niche construction dynamics. In nature, species do not live alone, and the most common mechanisms to prevent habitat degradation and population extinction are probably interactions among different species in a network. These interactions may open possibilities to new niches; negative effects for a species may be positive effects for others; and the complex feedback loops may maintain the stability of the whole network. This multidimensional aspect of niche construction is beyond the scope of the present article. Nevertheless, our study shows that rapid evolution, drift and intragenerational time lag in niche construction are important in delaying the spread of negative niche construction, therefore, it may buy more time for new species to come colonise and interact with the focal species and help establish a stable network. Lion et al. (2011) suggested that structured population, demographic and spatial alike, could favour the evolution of common goods and limit the spread of common “bads”. Our models suggest that a structure in time may contribute another dimension to the avoidance of such tragedies of commons. Here, time is particularly structured into population, niche construction and evolutionary dynamics, but it is not necessarily the only way. This further raises questions of how time could be structured in different ways.

## Acknowledgement

We would like to thank Florence Débarre, who also contributed largely to the conceiving of the project and who provided very helpful comments and advice during the execution of the project. This work was funded by the Chaire “Modélisation Mathématique et Biodiversité” of VEOLIA-Ecole Polytechnique-MNHN.

