## Supplementary for "Implications of drift and rapid evolution on negative niche construction"

### 1 Supplementary 1. Equilibrium of the unstructured population

The ecological dynamics using linear function for the waste production $f(c) = hc$ and additional mortality due to pollution $\omega(W) = vW$ have three equilibria. A nontrivial equilibrium where the population cannot survive

$$R_0^* = \frac{I_R}{\delta_R}$$
$$W_0^* = \frac{I_W}{\delta_W}$$
$$S_0^* = 0,$$

and two non zero equilibria

$$R_1^* = -\frac{\sqrt{4c\delta_W hI_R\rho v + (d\delta_W - \delta_R hv + I_W v)^2} - (d\delta_W - \delta_R hv + I_W v)}{2c\delta_W \rho} \tag{S.1}$$
$$R_2^* = \frac{\sqrt{4c\delta_W hI_R\rho v + (d\delta_W - \delta_R hv + I_W v)^2} + d\delta_W - \delta_R hv + I_W v}{2c\delta_W \rho} \tag{S.2}$$
$$W_i^* = \frac{hI_R}{\delta_W R_i^*} - \frac{\delta_R h - I_W}{\delta_W} \tag{S.3}$$
$$S_i^* = \frac{I_R - \delta_R R_i^*}{cR_i^*} \tag{S.4}$$

where $i = 1, 2$. The equilibrium with subscript 1 is not biologically meaningful because the resource density $R_1$ is always negative. We can indeed show that since all parameters are chosen positive, $4c\delta_W hI_R\rho v > 0$ and thus

$$\sqrt{4c\delta_W hI_R\rho v + (d\delta_W - \delta_R hv + I_W v)^2} > |d\delta_W - \delta_R hv + I_W v|$$

If $(d\delta_W - \delta_R hv + I_W v) > 0$ then $\sqrt{4c\delta_W hI_R\rho v + (d\delta_W - \delta_R hv + I_W v)^2} > (d\delta_W - \delta_R hv + I_W v)$

and $R_1^*$ is negative. If $(d\delta_W - \delta_R hv + I_W v) < 0$ then $\sqrt{4c\delta_W h I_R \rho v + (d\delta_W - \delta_R hv + I_W v)^2} - (d\delta_W - \delta_R hv + I_W v) > 0$ and $R_1^*$ is negative. Similarly, $R_2^* > 0$

Using Mathematica, it can be shown that $S_2^*$ and $W_2^*$ are positive when the consumption rate is sufficiently large and satisfies

$$c > \frac{d\delta_R\delta_W + \delta_R I_W v}{\delta_W I_R \rho} \tag{S.5}$$

#### Stability of the equilibrium

Equilibrium $(R_2^*, W_2^*, S_2^*)$ is stable if and only if all the eigenvalues of the jacobian matrix of the system at this point are negative. Here, the Jacobian matrix reads:

$$\begin{pmatrix} -cS - \delta_R & 0 & -cR \\ 0 & -\delta_W & ch \\ cS\rho & -Sv & -d - vW + cR\rho \end{pmatrix}$$

The eigenvalues $\lambda$ are the roots of the polynomial

$$\mathcal{C}_3\lambda^3 + \mathcal{C}_2\lambda^2 + \mathcal{C}_1\lambda + \mathcal{C}_0 = 0$$

where

$$\mathcal{C}_0 = (-cS_2^* - \delta_R)cS_2^* hv - cS_2^*(d + vW_2^*)\delta_W + \delta_R\delta_W(cR_2^*\rho - d - vW_2^*)$$

$$\mathcal{C}_1 = -cS_2^* hv - cS_2^*(d + vW_2^*) - cS_2^*\delta_W - \delta_R\delta_W + (cR_2^*\rho - d - vW_2^*)(\delta_R + \delta_W)$$

$$\mathcal{C}_2 = -cS_2^* - \delta_R - \delta_W + +cR_2^*\rho - d + vW_2^*$$

$$\mathcal{C}_3 = -1$$

are the coefficients of the polynomial. If we replace the equilibrium with its expression, we can write $\mathcal{C}_2$ as

$$\mathcal{C}_2 = -\frac{hv(\delta_R + 2\delta_W) + \delta_R hv + \mathcal{N}}{2hv}$$

where $\mathcal{N}$ is the numerator of the expression of $S_2^*$, which is always positive. Thus, $\mathcal{C}_2$ is always negative. Using Mathematica, we obtain that the coefficients $\mathcal{C}_0$ and $\mathcal{C}_1$ are negative. The polynomial thus has four negative coefficients, implying that there are two negative real roots at maximum, according to Descarte's rule. Furthermore, we consider the condition under which the roots of the polynomial have real negative parts to ensure the stability of the equilibrium. This happens when $\mathcal{C}_1\mathcal{C}_2 > \mathcal{C}_3\mathcal{C}_0$. After some algebra, we have

$$\mathcal{C}_1\mathcal{C}_2 - \mathcal{C}_3\mathcal{C}_0 = \left(\frac{cI_R\rho hv}{\delta_W} - (d\delta_W + I_W v) + hv(\delta_R + \delta_W + hv)\right)\delta_W\mathcal{N} + \\ (cI_R\rho\delta_W - \delta_R(d\delta_W + I_W v) + \delta_R hv(\delta_R + \delta_W))2\delta_W hv$$

Due to condition (S.5), we have $cI_R\rho\delta_W - \delta_R(d\delta_W + I_W v) > 0$. The above expression is negative when $hv$ is extremely small, which implies that the rate of waste production is small and/or the mortality rate due to pollution is small. In such a case, the population of consumer will likely tend to infinity because it is is mainly limited by the waste in the environment. When there is little waste due to low waste production or when the population is not vulnerable to the pollution, the equilibrium become unstable and increase to infinity.

### 2 Supplementary. Invasion fitness and basic reproduction ratio of mutant of the unstructured population dynamics

The invasion fitness of a mutant that adopts a different consumption rate value is simply its per capital growth rate evaluated at the equilibrium of the resident $(R^*, W^*)$ since the model is linear, and it can be written as

$$\frac{1}{S_m}\frac{dS_m}{dt} = c_m\rho R^*(c) - d - vW^*(c)$$

The mutant can invade if its invasion fitness is positive, which, after some algebra, can be written as

$$\frac{c_m\rho R^*(c)}{d + W^*(c)} > 1$$

The left-hand side of the above inequality is the basic reproduction ratio of the mutant $F(c_{mut}, c)$ in the main text.

The selection gradient of the consumption rate is

$$\frac{\partial \frac{1}{S_m}\frac{dS_m}{dt}}{\partial c_m} = \rho R^*(c)$$

which is always positive.

### 3 Supplementary. Equilibrium of the structured population without resources dynamic

System (3) in the main text has three equilibria. A nontrivial equilibrium where the population cannot survive

$$A_0^* = J_0^* = 0$$
$$W_0^* = \frac{I_W}{\delta_W}$$

and two non zero equilibria

$$A_1^* = \frac{X+Y}{Z}$$
$$A_2^* = \frac{X-Y}{Z}$$
$$J_i^* = \frac{A_i^*(d_A + v_A W_i^*)}{c_J R}$$
$$W_1^* = -\frac{K + (c_J R + d_J)v_A + d_A v_J}{2 v_A v_J}$$
$$W_2^* = \frac{K - v_A(c_J R + d_J) - d_A v_J}{2 v_A v_J}$$

where $i = 1, 2$

$$X = c_J R v_A(\delta_W p_A R - h_J(\delta_W(d_A - 2rR) + I_W v_A)) - d_J v_A(d_A \delta_W h_J + h_J I_W v_A - \delta_W p_A R) + I_W v_A v_J(d_A h_J + 2p_A R) + d_A \delta_W v_J(d_A h_J + p_A R)$$

$$Y = d_A \delta_W h_J + h_J I_W v_A + \delta_W p_A R K$$

$$Z = 2v_A(c_J h_J R v_A(h_J r + p_A) - p_A v_J(d_A h_J + p_A R) + d_J h_J p_A v_A)$$

$$K = \sqrt{(c_J R v_A - d_A v_J + d_J v_A)^2 + 4c_J r R^2 v_A v_J}$$

The equilibrium $(A_1^*, J_1^*, W_1^*)$ is not biologically meaningful because the waste density $W_1^*$ is always negative because $K$ is a square root and other demographic parameters are positive.

Using Mathematica, we found that equilibrium $(A_2^*, J_2^*, W_2^*)$ is positive when

$$\frac{c_J R \rho R}{\mathcal{D}_J \mathcal{D}_A} > 1 \tag{S.6}$$

where

$$\mathcal{D}_A = d_A + v_A W_0^* \tag{S.7}$$

$$\mathcal{D}_J = c_J R + d_J + v_J W_0^* \tag{S.8}$$

The left hand side of inequality (S.6) is the basic reproduction ratio of the consumer (expression (4) in the main text).

### 4 Supplementary. Stability of equilibrium

The stability of the positive equilibrium $A_2^*, J_2^*, W_2^*$ in section 3 is determined by the sign of the eigenvalues of the jacobian matrix of system (3) in the main text is

$$\begin{pmatrix} -d_A - v_A W & c_J & -A v_A \\ r & -c_J - d_J - v_J W & -J v_J \\ p_A & c_J h_J & -\delta_W \end{pmatrix}$$

The eigenvalues $\lambda$ of the above jacobian matrix are the roots of the following polynomial

$$C_3\lambda^3 + C_2\lambda^2 + C_1\lambda + C_0$$

where

$$C_3 = -1$$

$$C_2 = -D_A - D_J - \delta_W$$

$$C_1 = -D_A D_J + c_J c_A R^2 - A_2^* p_A v_A - c_J h_J v_J J_2^* - \delta_W (D_A + D_J)$$

$$C_0 = (-D_A D_J + c_J c_A R^2)\delta_W - A_2^* v_A (c_J h_J r R + D_J p_A) - c_J v_J (D_A h_J + p_A R) J_2^*$$

where $D_A$ and $D_J$ are as in expression (S.7). Coefficients $C_3$ and $C_2$ are always negative. When the population is at equilibrium $(W_2^*, J_2^*, A_2^*)$, we have $-D_A D_J + c_J c_A R^2 = 0$, thus $C_1$ and $C_0$ are also negative because the equilibrium is positive and other demographic parameters are also positive.

As a result the polynomial always has four negative coefficients, suggesting that it can have at maximum two negative real roots according to Descarte's rule. This also make the first condition of the Routh-Hurwitz satisfied. However, this condition is insufficient for stable $J_2^*, A_2^*$, and $W_2^*$ because the polynomial can have imaginary roots with positive real part. Under the second Routh-Hurwitz condition, the roots of the polynomial have their real part negative if $C_1 C_2 > C_3 C_0$ which leads to the following condition

$$X_1 + X_2 > 0 \tag{S.9}$$

where

$$X_1 = \delta_W(D_A + D_J)(D_A + D_J + \delta_W)$$

$$X_2 = A_2^*((v_A - Rv_J)(D_A p_A - c_J h_J c_A R) + \delta_W(D_A h_J v_J + p_A v_A))$$

where $D_A$ and $D_J$ are again as in expression (S.7). We observe that $X_1$ is always positive, but $X_2$ can be negative or positive. We present the possible signs for $X_1 + X_2$ in the following table

| Conditions | $X_2$ | $X_1 + X_2$ | Status of $J_2^*, A_2^*, W_2^*$ |
|---|---|---|---|
| $\frac{v_A}{v_J} = R$ | + | + | +, stable |
| $\frac{v_A}{v_J} < R$<br>$\delta_W < D_A\left(\frac{v_J}{v_A}R - 1\right)$<br>$h_J > -\frac{p_A(D_A v_J(\frac{v_A}{v_J} - R) + \delta_W v_A)}{c_J c_A R v_J(R - \frac{v_A}{v_J}) + D_A \delta_W v_J}$ | + | + | +, stable |
| $\frac{v_A}{v_J} < R$<br>$\delta_W < D_A\left(\frac{v_J}{v_A}R - 1\right)$ | + | + | +, stable |
| $\frac{v_A}{v_J} > R$<br>$\delta_W < \frac{c_J c_A R(\frac{v_A}{v_J} - R)}{D_A}$<br>$h_J < -\frac{p_A(D_A v_J(\frac{v_A}{v_J} - R) + \delta_W v_A)}{c_J c_A R v_J(R - \frac{v_A}{v_J}) + D_A \delta_W v_J}$ | + | + | +, stable |
| $\frac{v_A}{v_J} > R$<br>$\delta_W \geq \frac{c_J c_A R(\frac{v_A}{v_J} - R)}{D_A}$ | + | + | +, stable |
| $\frac{v_A}{v_J} > R$<br>$\delta_W < \frac{c_J c_A R(\frac{v_A}{v_J} - R)}{D_A}$<br>$h_J > -\frac{p_A(D_A v_J(\frac{v_A}{v_J} - R) + \delta_W v_A)}{c_J c_A R v_J(R - \frac{v_A}{v_J}) + D_A \delta_W v_J}$ | - | not clear | not clear |
| $\frac{v_A}{v_J} < R$<br>$\delta_W < D_A\left(\frac{v_J}{v_A}R - 1\right)$<br>$h_J < -\frac{p_A(D_A v_J(\frac{v_A}{v_J} - R) + \delta_W v_A)}{c_J c_A R v_J(R - \frac{v_A}{v_J}) + D_A \delta_W v_J}$ | - | not clear | not clear |

Table 1: Conditions for the sign of $X_2$, $X_1 + X_2$ and the stability of the equilibrium

The main conclusion from Table 1 is that for most of the parameter ranges, the positive equilibrium $(A_2^*, J_2^*, W_2^*)$ is stable. The equilibrium may be unstable when $X_2$ is negative, but unless $X_2$ is more negative than $X_1$ the equilibrium will be stable. In fact, in a numerical analysis, we show that considering a range of the $h_J$ value, $X_2$ is negative but the whole expression $X_1 + X_2$ is always positive, hence this equilibrium is always positive and stable (Figure S. 1).

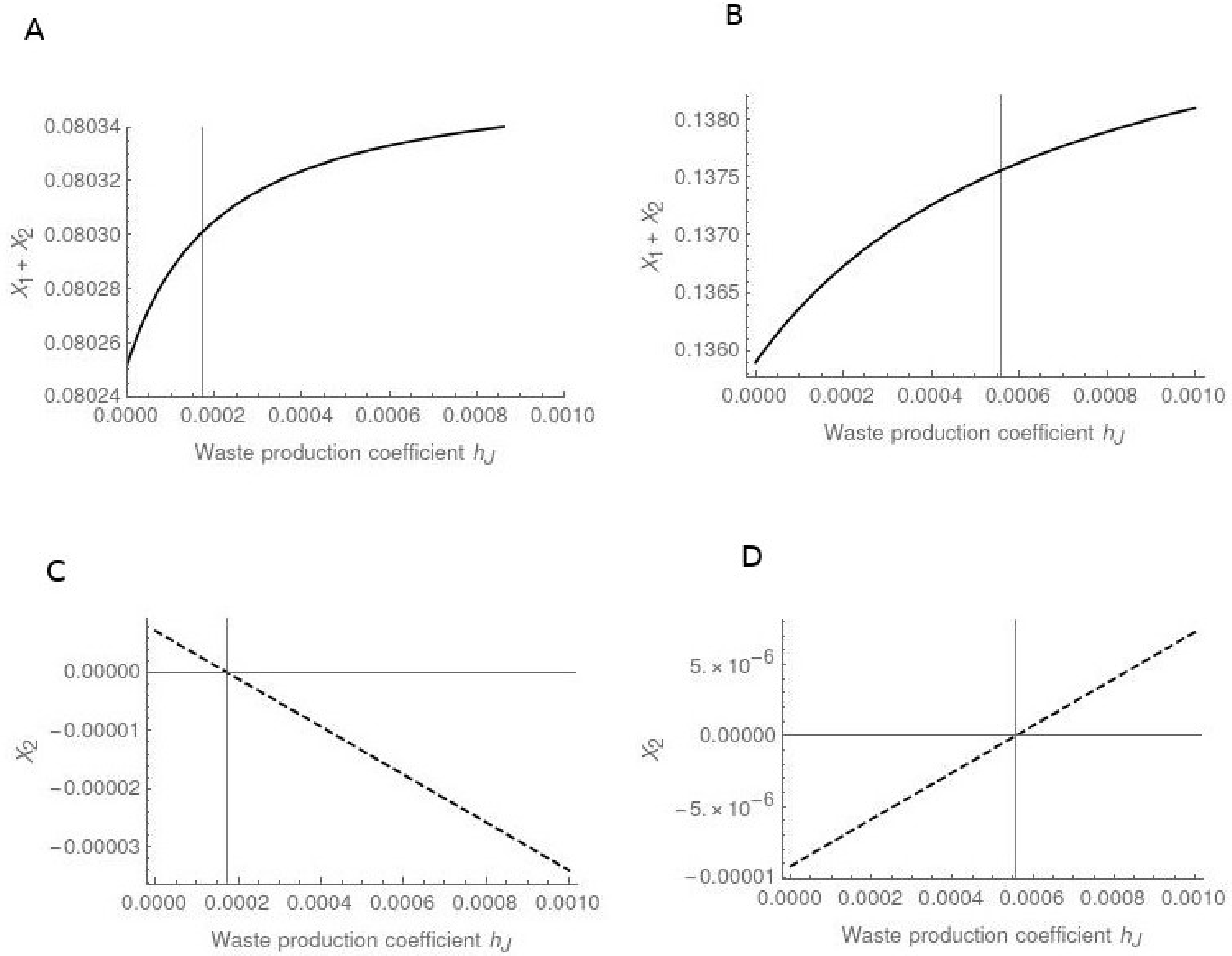


Figure S. 1: $X_1 + X_2$ is always positive while $X_2$ can be negative depending on the threshold of $h_J$ (gray vertical lines) in the last two conditions in Table 1. Common parameters $d_J = 0.00011, d_A = 0.1, v_A = 0.008, v_J = 0.01, I_W = 1.3, p_A = 0.001, c_J = 0.1, \delta_W = 0.008$. A, C) $R = 0.3, \rho = 275.1$, B, D) $R = 1.3, r = 25.1$

### 5 Supplementary. Invasion fitness of the mutant

The dynamics of a mutant that arises in the resident population with dynamics (3) are

$$\frac{dJ_m}{dt} = \rho R A_m - d_J J_m - v_J W^* J_m - c_{Jm} R J_m \tag{S.10a}$$

$$\frac{dA_m}{dt} = c_{Jm} R J_m - d_A A_m - v_A W^* A_m \tag{S.10b}$$

where $W^*$ is the density of the waste at equilibrium. Here, only equilibrium 2 in the Supplementary document 2 is qualified to use, hence $W^* = W_2^*$. The Jacobian matrix of system (S.10) is

$$J_{inv} = \begin{pmatrix} -d_A - v_A W_2^* & c_{Jm} R \\ rR & -c_{Jm} R - d_J - v_J W_2^* \end{pmatrix} \tag{S.11}$$

A mutant can invade if at least one eigenvalue of matrix (S.11) is positive. An equivalent of this condition is that either the determinant of matrix (S.11) is negative, indicating that there is one positive and one negative eigenvalues, or both the determinant and trace of matrix (S.11) are positive. However, here the trace of matrix (S.11) is always negative. Therefore, the only condition that can be satisfied is that the matrix's determinant is negative, which leads to

$$\frac{c_{Jm} r R^2}{D_A D_J(c_{Jm})} > 1$$

where

$$D_A = d_A + v_A W_2^*$$

$$D_J(c_{Jm}) = d_J + c_{Jm} R + v_J W_2^*$$

The left-hand side of the above inequality is the basic reproduction ratio $F_{mut}$ of a mutant in the main text (expression (5)).

#### Relationship between the selection gradient and the derivative of the determinant of $J_{inv}$

In this section, we will show that the sign of the selection gradient is opposite to the sign of the derivative of the determinant of $J_{inv}$ for most of the cases. Therefore, the determinant can serve as an approximation for the invasion fitness but the conditions that select for lower growth rate must be derived from the selection gradient.

The eigenvalues of matrix (S.11) are

$$\lambda_1 = \frac{1}{2}\left(\sqrt{4c_{Jm}rR^2 + (D_A(W_2^*) - D_J(W_2^*))^2} - (D_A(W_2^*) + D_J(W_2^*))\right) \tag{S.12}$$

$$\lambda_2 = \frac{1}{2}\left(-\sqrt{4c_{Jm}rR^2 + (D_A(W_2^*) - D_J(W_2^*))^2} - (D_A(W_2^*) + D_J(W_2^*))\right) \tag{S.13}$$

where $\lambda_1$ is the leading eigenvalue, $\lambda_2$ is always negative, and $|\lambda_1| < |\lambda_2|$. The selection gradient of the trait value is

$$\frac{\partial \lambda_1}{\partial c_{Jm}} = \frac{1}{2}R\left(\frac{2rR - (D_A(W_2^*) - D_J(W_2^*))}{\sqrt{4c_{Jm}rR^2 + (D_A(W_2^*) - D_J(W_2^*))^2}} - 1\right)$$

which will always be positive when the mutant trait value is small as $4c_{Jm}rR^2 \approx 0$. When the trait value is large, the sign of the selection gradient depends on both the mutant trait value and the waste concentration in the environment. Using Mathematica, the solution for negative selection gradient, which implies selection for lower growth rate, is

$$\begin{cases} v_A > v_J & \text{(S.14)} \\ W^* > \frac{d_J + rR - d_A}{v_A - v_J}\Big|_{c_J = c_{Jm}} . & \text{(S.15)} \end{cases}$$

This is the condition (6) in the main text.

The derivative of the determinant of $J_{inv}$ with respect to $c_{Jm}$ is

$$\frac{\partial Det(J_{inv})}{\partial c_{Jm}} = \lambda_2 \frac{\partial \lambda_1}{\partial c_{Jm}} + \lambda_1 \frac{\partial \lambda_2}{\partial c_{Jm}} = R(-rR + D_A(W_2^*))$$

We have

$$\frac{\partial \lambda_2}{\partial c_{Jm}} = -\frac{1}{2} R \left( \frac{2rR - (D_A(W_2^*) - D_J(W_2^*))}{\sqrt{4c_{Jm} rR^2 + (D_A(W_2^*) - D_J(W_2^*))^2}} + 1 \right)$$

Therefore it can be easily seen that if $\partial\lambda_1/\partial c_{Jm} > 0$ then $\partial\lambda_2/\partial c_{Jm} < 0$. Moreover, $\lambda_2 < 0$ and $\lambda_1 > 0$ for when the invasion condition is satisfied. Thus $\lambda_2 \partial\lambda_1/\partial c_{Jm} < 0$ and $\lambda_1 \partial\lambda_2/\partial c_{Jm} < 0$, hence $\partial Det(J_{inv})/\partial c_{Jm} < 0$.

If $\partial\lambda_1/\partial c_{Jm} < 0$, then $\lambda_2 \partial\lambda_1/\partial c_{Jm} > 0$. If $\partial\lambda_2/\partial c_{Jm} > 0$, then $\lambda_1 \partial\lambda_2/\partial c_{Jm} > 0$, and $\partial Det(J_{inv})/\partial c_{Jm} > 0$. If $\partial\lambda_2/\partial c_{Jm} < 0$, then the sign of $\partial Det(J_{mut})/\partial c_{Jm}$ is undecided.

We can summarise the signs of different term and conclude the sign of the selection gradient and $\partial Det(J_{inv})/\partial c_{Jm}$ in the following table

| $\lambda_1$ | $\lambda_2$ | $\partial\lambda_1/\partial c_{Jm}$ (Selection gradient) | $\partial\lambda_2/\partial c_{Jm}$ | $\partial Det(J_{inv})/\partial c_{Jm}$ |
|---|---|---|---|---|
| + | - | + | - | - |
| + | - | - | + | + |
| + | - | - | - | undecided |

### 6 Supplementary: Effect of growth rate value on the waste density at equilibrium

The derivative of the waste value at equilibrium with respect to the growth rate is

$$\frac{dW_2^*}{dc_J} = \frac{R}{2v_J}\left(\frac{Q + 2rRv_J}{\sqrt{4c_J rR^2 v_A v_J + Q^2}} - 1\right) \tag{S.16}$$

where $Q = (dJ + cJR)vA - dAvJ$. Expression (S.16) is positive if

$$Q + 2rRv_J > \sqrt{4c_J rR^2 v_A v_J + Q^2} \tag{S.17}$$

It can be seen that $Q$ is always positive because of condition (4). We can square both sides of the above expression and obtain

$$4rRvJ(dJvA - dAvJ + rRvJ) > 0$$

which is always satisfied becausse of condition (4). Alternatively, using the function Reduce in Mathematica with assumption that all demographic parameters are positive and the condition for positive equilibrium has to be satisfied, we can also prove that expression (S.16) is always positive.

### 7 Supplementary Figure

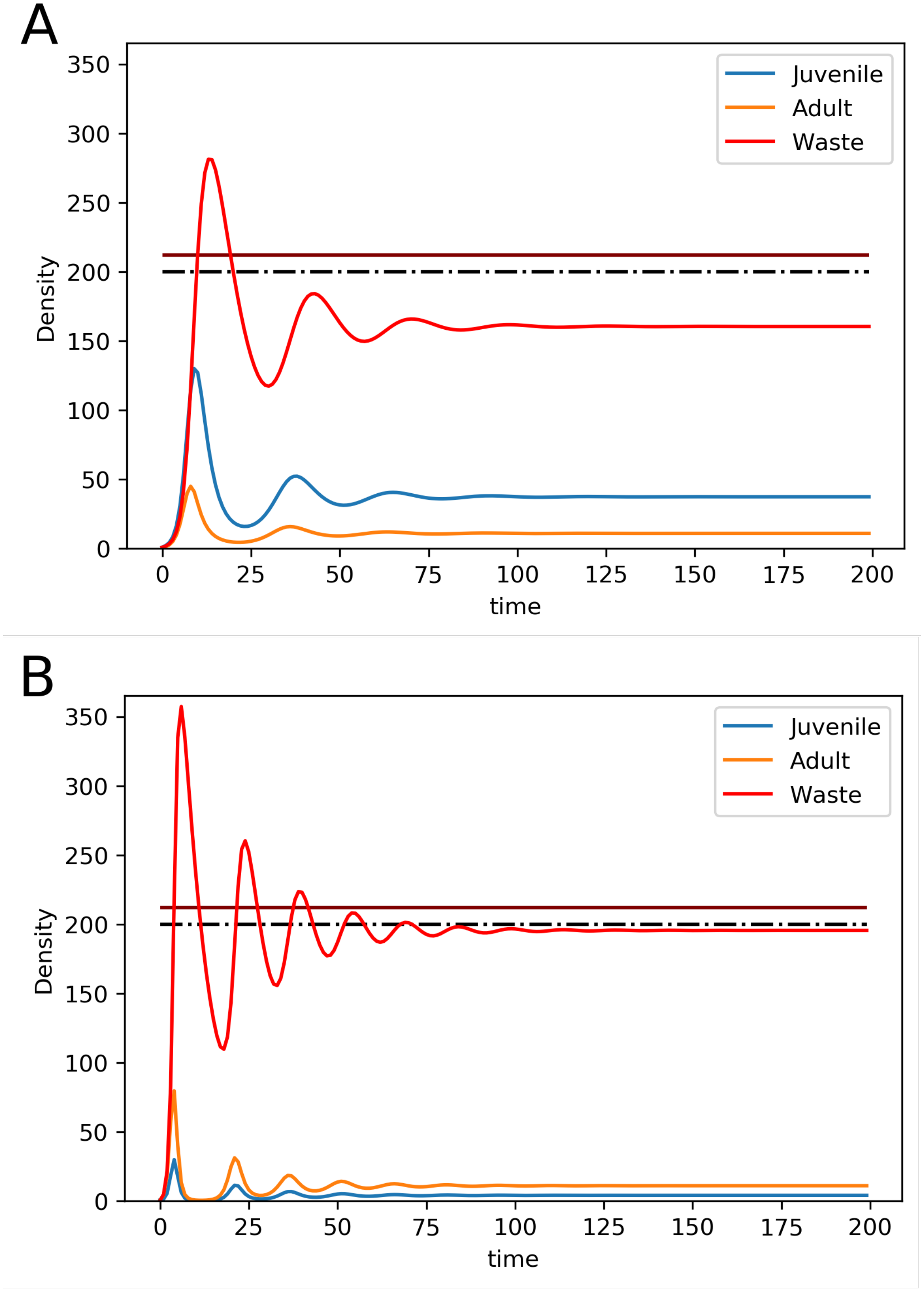


Figure S. 2: Dynamics of a resident population and waste concentration. The dark red horizontal line indicates the threshold above which selection will favour mutants with smaller growth rate (The expression of the line is $(rR - d_A - d_J)/(v_A - v_J)$). The dash dot line indicates the value to which the waste density at equilibrium is asymptotic as the growth rate value increases (The expression of the line is $(rR - d_A)/v_A$)). A) Resident population with small growth rate $c_J = 0.5$, B) Resident population with larger growth rate $c_J = 5.5$. Other parameters: $d_J = 0.1, d_A = 0.1, h_J = 1.1, v_J = 0.0001, v_A = 0.01, I_W = 0.3, \delta_W = 0.13, c_A = 2.1, p_A = 0.001, R = 1$

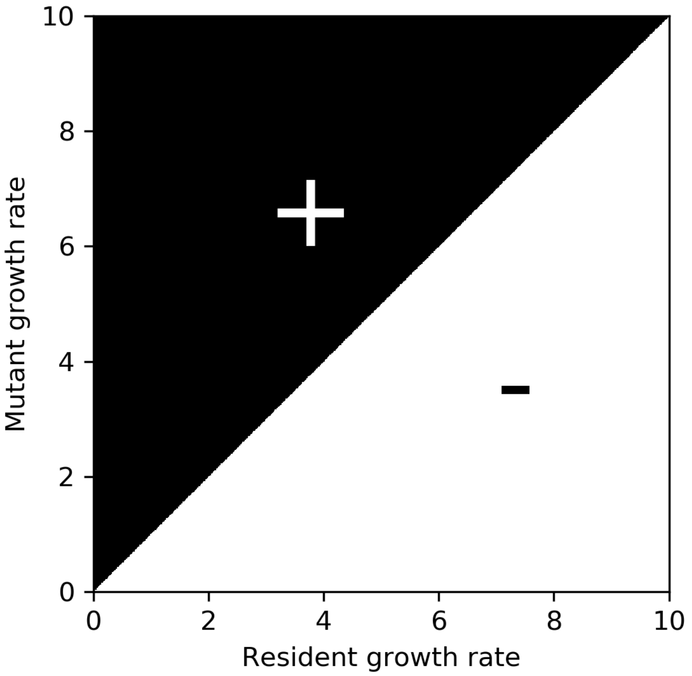


Figure S. 3: Pairwise invisibility plot showing that higher growth rate is always favoured. The black area indicates the area where mutants can invade, which is annotated by the plus sign. The white area indicates the area where mutants cannot invade, which is annotated by the minus sign. Parameters: $d_J = 0.1, d_A = 0.1, h_J = 1.1, v_J = 0.0001, v_A = 0.01, I_W = 0.3, \delta_W = 0.13, c_A = 2.1, p_A = 0.001$

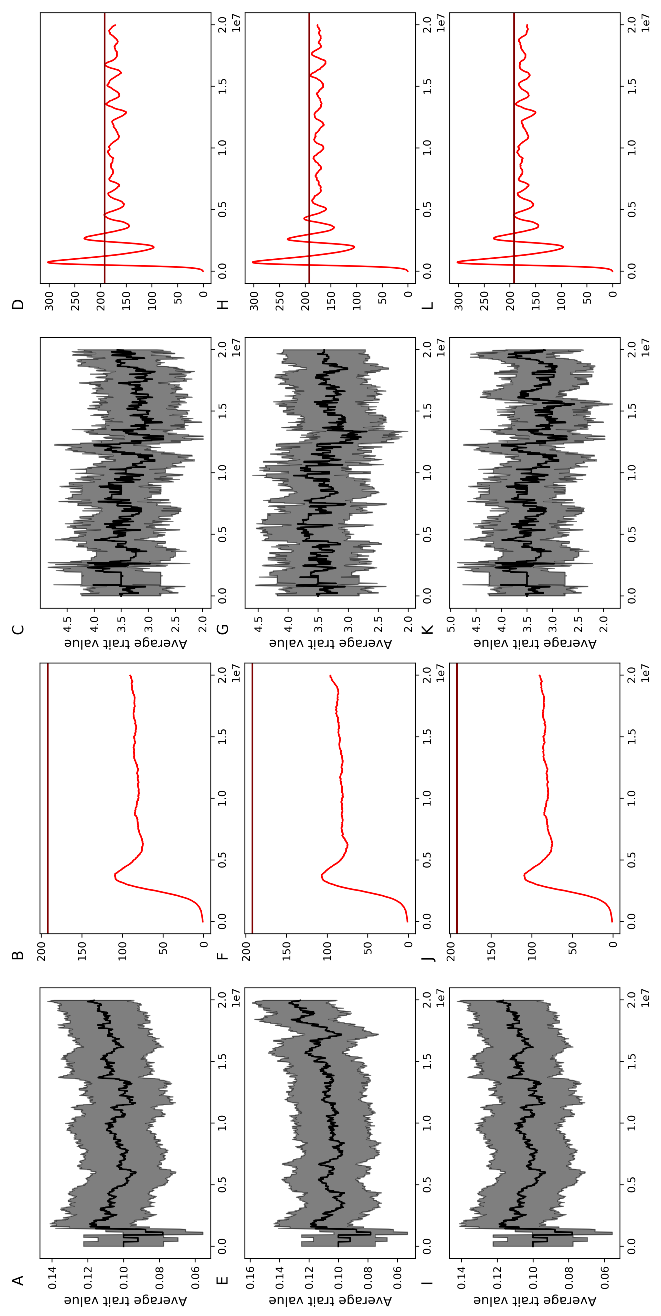

Figure S. 4: A, B, E, F, I, J. Different replicates of simulations where populations start with extremely slow growth rate. Parameters are the same as those in the right panels of Figure 4 in the main text. B, C, G, H, K, L. Different replicates of simulations where populations start with higher growth rate. Gray areas indicate the standard deviation of the average trait value. Parameters are the same as those in the right panels of Figure 3.

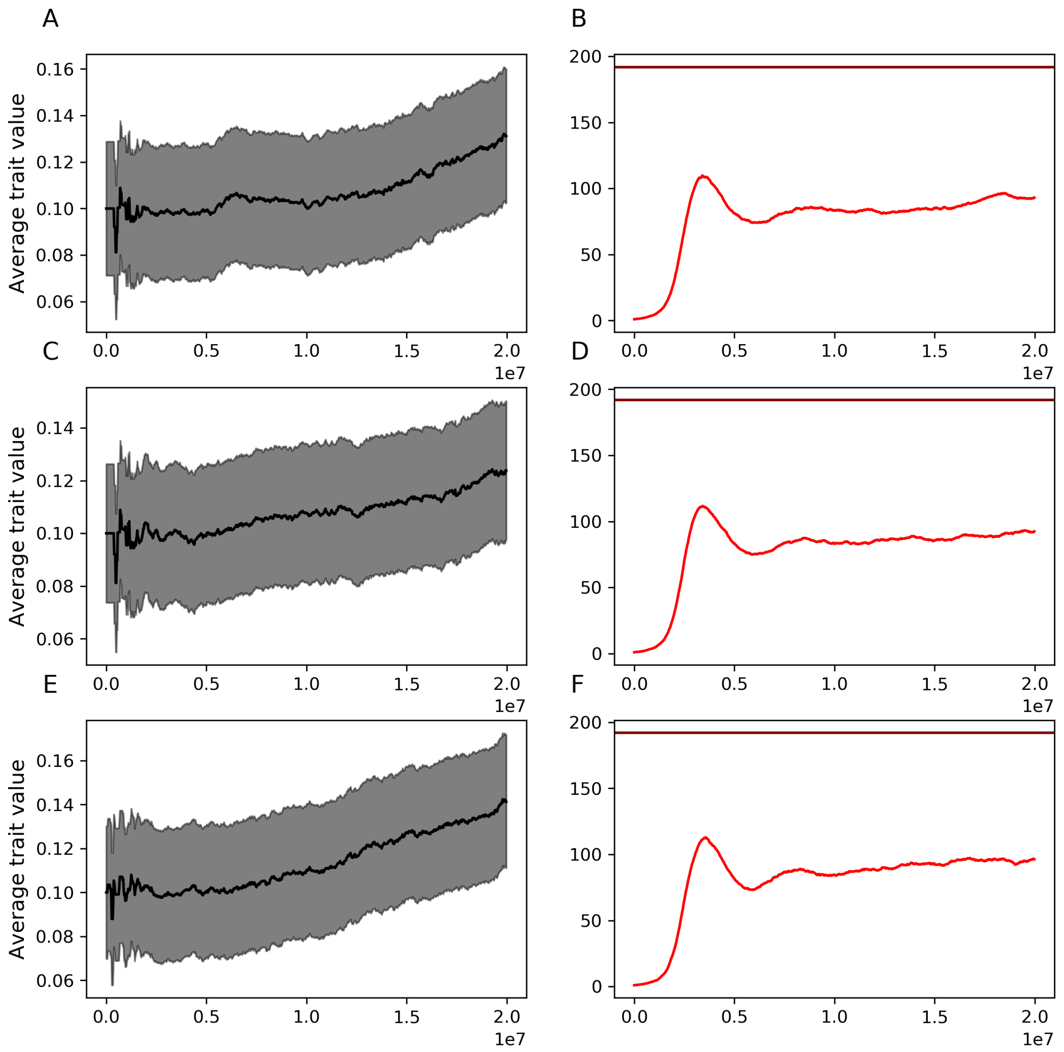


Figure S. 5: Each row is a different replicate of the simulation where mutation rate is high $m = 0.01$ and the population and waste dynamics are slow. Gray areas indicate the standard deviation of the average trait value. Other parameters are the same as those in the right panels of Figure 4

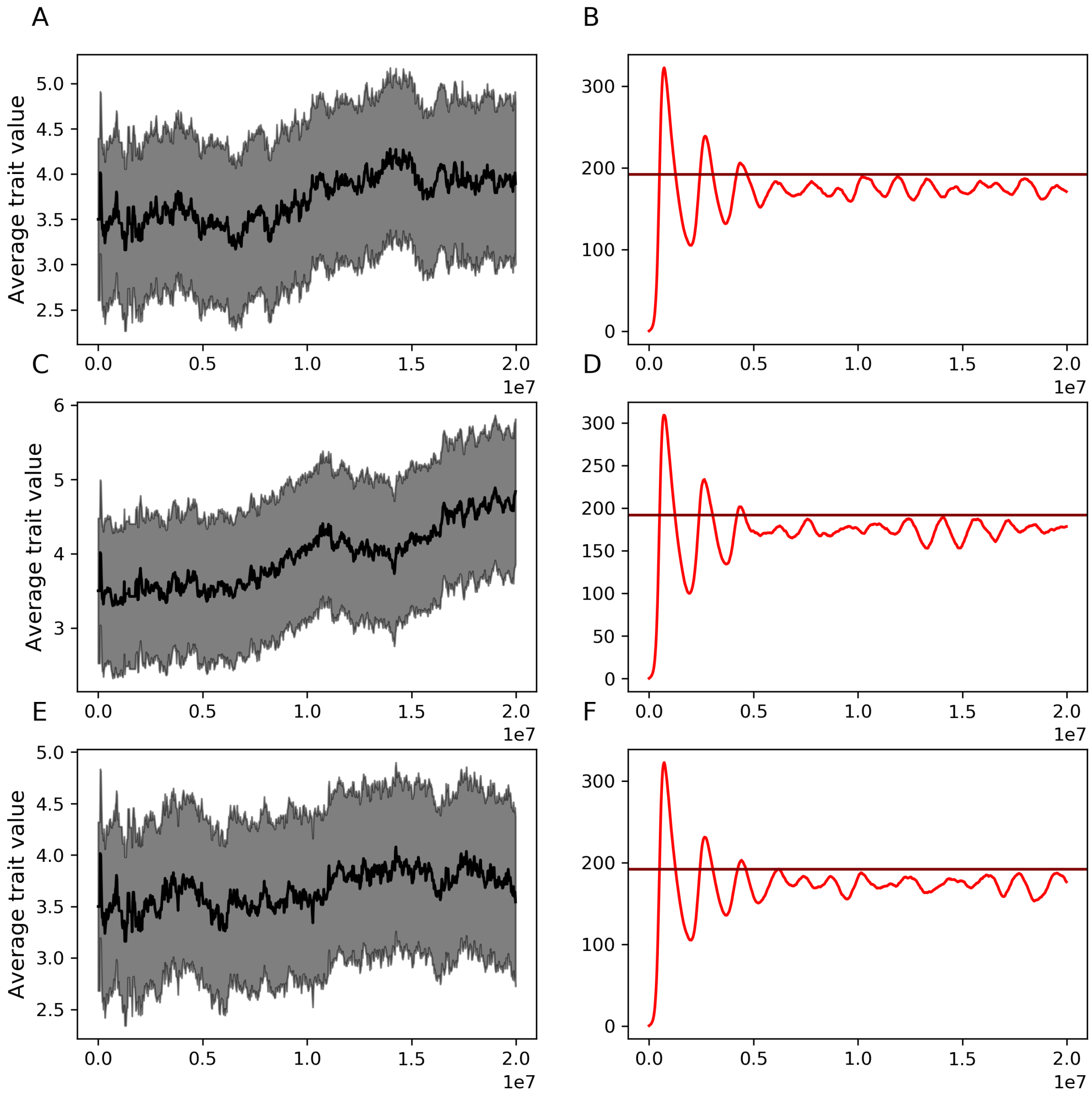


Figure S. 6: Each row is a different replicate of the simulation where mutation rate is high $m = 0.01$ and the population and waste dynamics are slow. Other parameters are the same as those in the right panels of Figure 4

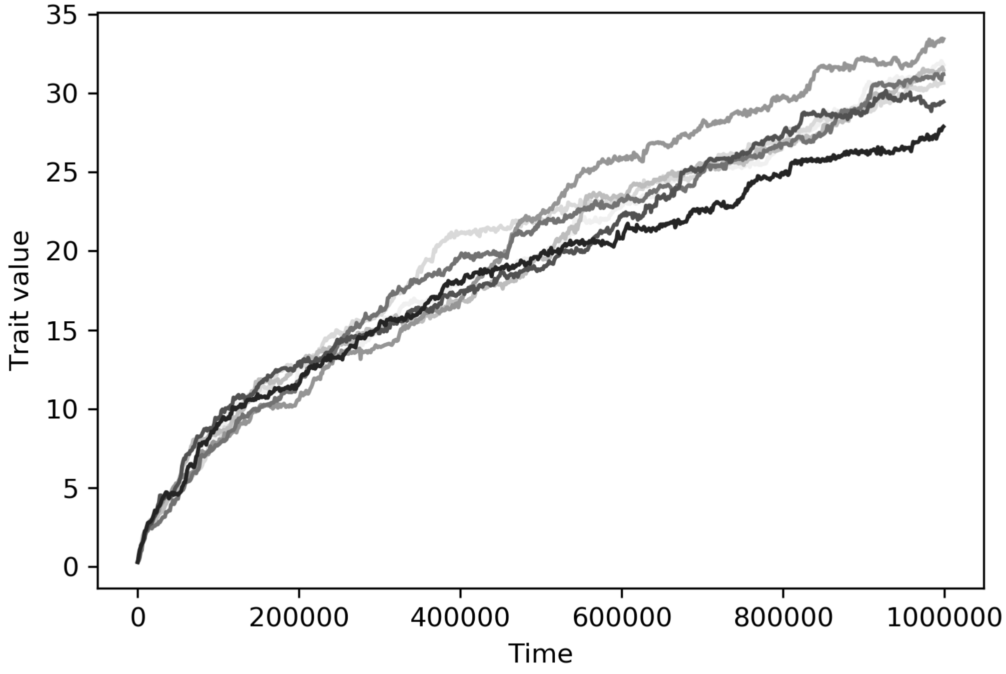


Figure S. 7: Simulations with different sizes of waste pool. Small pool: $I_W = 0.01024, \delta_W = 0.0003$, intermediate pool: $I_W = 3, \delta_W = 0.3$, big pool: $I_W = 1024, \delta_W = 30$. Other parameters are the same as in Figure 5.
